# The influence of sponge-dwelling gobies (*Elacatinus horsti*) on feeding efficiency of Caribbean sponge hosts, *Aplysina lacunosa* and *Aplysina archeri*

**DOI:** 10.1101/2021.11.30.470666

**Authors:** Megan J. Siemann, Aldo Turco

## Abstract

Mutualistic associations between benthic marine invertebrates and reef taxa are common. Sponge-dwelling gobies benefit from protection within sponge tubes and greater food availability. Sponge-dwelling gobies are hypothesized to increase sponge pump rates by consuming polychaete parasites, but such increases have not yet been demonstrated. We investigated the association between sponge-dwelling gobies (*Elacatinus horsti*) and two species of tube sponge (*Aplysina lacunosa* and *Aplysina archeri*) in Bonaire, Caribbean Netherlands. We visually assessed goby presence in sponges and used *in situ* methods with fluorescein dye to estimate feeding rates via pump rates. *Aplysina archeri* were more likely to host a goby than *A. lacunosa*. For both sponge species, pump rates of tubes with gobies were higher on average than those of tubes without gobies. Our observations, therefore, suggest that *E. horsti* associations with *Aplysina* are likely mutualistic relationships in which sponges benefit from higher feeding rates when gobies are present.

## Introduction

Being active suspension feeders, sponges are recognized as important planktonic feeders in coral reef food webs (Lesser 2006) with the potential to alter the chemical and biological environment of the water column. Investigating the magnitude of sponge filtration and the factors that influence pumping rates and feeding efficiency provides a basis for understanding how the role of sponges might impact local ecosystems. For instance, at high biomasses, sponges reduce picoplankton availability of the water column which impacts food availability for planktivores (Savarese et al. 1997). Water pumping rates through the aquiferous system of sponges has been widely accepted as the common measure of their feeding efficiency (Gerrodette and Flechsig 1979; Hendler 1984; Savarese et al. 1997; McMurray et. al 2014; Morganti et al. 2019). This is influenced by a number of organisms that interact with sponges; oysters and brittle stars, for example, have a symbiotic relationship with sponges in which the sponge benefits from increased pump rates (Hendler 1984; Burns and Bingham 2002). Reviews have categorized interspecific interactions of sponges as predation, competition, or utilization of sponges as microhabitat (Rocha et al. 2000; Wulff 2006; Bell 2008). For instance, a large number of invertebrates and fish species dwell inside sponge microhabitats (Wulff 2006; Bell 2008).

Gobies associated with sponges include free-swimming and obligate sponge-dwelling species (Tyler and Böhlke 1972; Colin 1975; Randall and Lobel 2009; D’Aloia et al. 2011). *Elacatinus horsti* (yellowline goby) is a sponge-dwelling goby that occupies convoluted barrel (*Aplysina lacunosa*) and stove-pipe sponges (*Aplysina archeri*), which are found in the Caribbean (Villamizar and Laughlin 1991). Both *A. archeri* and *A. lacunosa* grow in clumps of tubes with individual tubes reaching maximum lengths of 180 cm and 90 cm, respectively (Humann and DeLoach 1992).

While a variety of different organisms inhabiting sponges have been documented (Tyler and Böhlke 1972; Villamizar and Laughlin 1991; Meroz and Ilan 1995; Bell 2008; D’Aloia et al. 2011), the effects of the inhabitants on the sheltering sponge remain understudied. Hendler (1984) found that *Callyspongia vaginalis* sponges benefit from resident brittle stars that clear sediment from their tubes. The relationship between sponge-dwelling gobies and their *Evermannichthys* sponge hosts was originally described as an obligate commensalism with gobies benefitting from the shelter (Böhlke and Robins 1969). Other research has recognized sponge-dwelling invertebrates as an additional benefit to *Elacatinus* gobies. Colin (1975) concluded that *E. horsti* were eating parasitic polychaetes because sponge tissue was found in their guts.

Furthermore, it has been proposed that sponges with sponge-dwelling gobies benefit from lower parasite loads (Whiteman and Côté 2002; D’Aloia et al. 2011), which may increase sponge feeding efficiency. Directly measuring a sponges’ feeding efficiency *in situ* is considered difficult (Yahel et al. 2005). Therefore, pump rate has been considered a proxy for feeding efficiency on the assumption that the more water a sponge can pump, the more food can be filtered (Gerrodette and Flechsig 1979; Hendler 1984; Savarese et al. 1997; McMurray et. al 2014; Morganti et al. 2019). The pump rate of a species is also affected by the size of sponges and depth, which must be considered potential sources of variation (Lesser 2006; McMurray et al. 2014), and therefore, accounted for in studies on the feeding rates of sponge.

In this study, we quantified the effect of *E. horsti* presence on *Aplysina* spp. using pump rate as a proxy for feeding rate. We predicted that tubes of *Aplysina lacunosa* and *Aplysina archeri* containing *E. horsti* gobies would have a higher pump rates (*i*.*e*., greater feeding efficiency).

## Methods

### Study site

We conducted this study from September - October 2017 at Playa Lechi dive site (12°09’36.6”N, 68°16’55.1”W) on the leeward side of the island of Bonaire, Caribbean Netherlands. The study area is characterized by a reef slope starting at 5 m depth and continuing until ∼35 m to a flat sandy substrate. While *A. archeri* and *A. lacunosa* are found in depths of 3-45 m (Villamizar and Laughlin 1991), we conducted this SCUBA study between 5-15 m deep to limit depth-dependent effects on pump rate (Bell 2008). A three-week study duration and daytime dive times were chosen to limit seasonal and daily temporal cycles in sponge pump rates (Morganti et al. 2019).

### Investigation of sponge abundance and symbiont presence

We surveyed three 50 m transects to investigate the abundance of both *Aplysina* species and the frequency of *E. horsti*. Only presence and absence information was recorded for the gobies. To limit the effect of the divers, a 3-min acclimation period was allotted after laying the transect.

### Measuring pump rate

#### In situ methods

We randomly selected *A. archeri* and *A. lacunosa* and checked tubes for *E. horsti*. At each sponge, we measured tube length and pump rate on the most landward tube with *E. horsti* and, where possible, the most landward tube without *E. horsti*. We measured pump rates following the approach of Gerrodette and Flechsig (1979). We taped a flexible, clear tube (1 cm diameter) with scale marks to a hole in a 700 ml resealable, plastic bag. Then, we secured the bag to the sponge osculum using rubber bands. One diver injected 2 ml of 100 mg L^-1^ fluorescein dyed seawater (Yahel et al. 2005) through a hole near the base of the tube, while the other stabilized the tube and filmed dye movement at 30 frames s^-1^. For each sponge tube surveyed, we released dye three times then removed the bag.

#### Video processing

We used linear regressions to calculate the velocity of the dye front on a fixed 2D plane based on scale marks on the tube and frame rate (Tracker 4.11.0 software) and used the tube’s cross-section area to convert these to pump rates.

### Statistical analysis

We performed separate repeated measures (multiple measurements per tube) ANCOVAs for each species of sponge to determine whether pump rate depended on the presence of *E. horsti* (categorical predictor) and tube length (continuous predictor). *Elacatinus horsti* presence was treated as a binary variable as only 3 of 72 tubes tested for pump rate contained more than one *E. horsti*. We used a z-test (two-tail) to test for differences between the proportions of tubes containing *E. horsti* for the two species of sponge.

## Results and discussion

### Mutualistic symbiosis

Pump rates of both sponge species increased when *E. horsti* were present (*A. lacunosa*: F_1,32_=10.56, p=0.0027, +2.35 cm^3^ s^-1^; *A. archeri*: F_1,34_=8.39, p=0.0458, +1.78 cm^3^ s^-1^; Fig. 1). The significantly higher pump rates of tubes with *E. horsti* supported the hypothesis of a conferred feeding benefit to sponges. In nature, the symbiosis is thought to be obligatory for the sponge-dwelling goby that is dependent on the sponge for both food and shelter (Colin 1975; Whiteman and Côté 2004a; D’Aloia et al. 2011). *Elacatinus horsti* benefit from the interaction through nutrition supplied from eating parasites that feed on sponge tissue plus gobies are readily attacked by predators when disturbed from their sponge tubes or held in open, experimental tanks (Colin 1975). While our results are consistent with a benefit to the sponge and therefore a mutually beneficial symbiosis, it may also be that gobies choose tubes with higher pump rates. Such a possibility would require experimental introductions or removals of gobies to be tested.

**Fig. 1.**
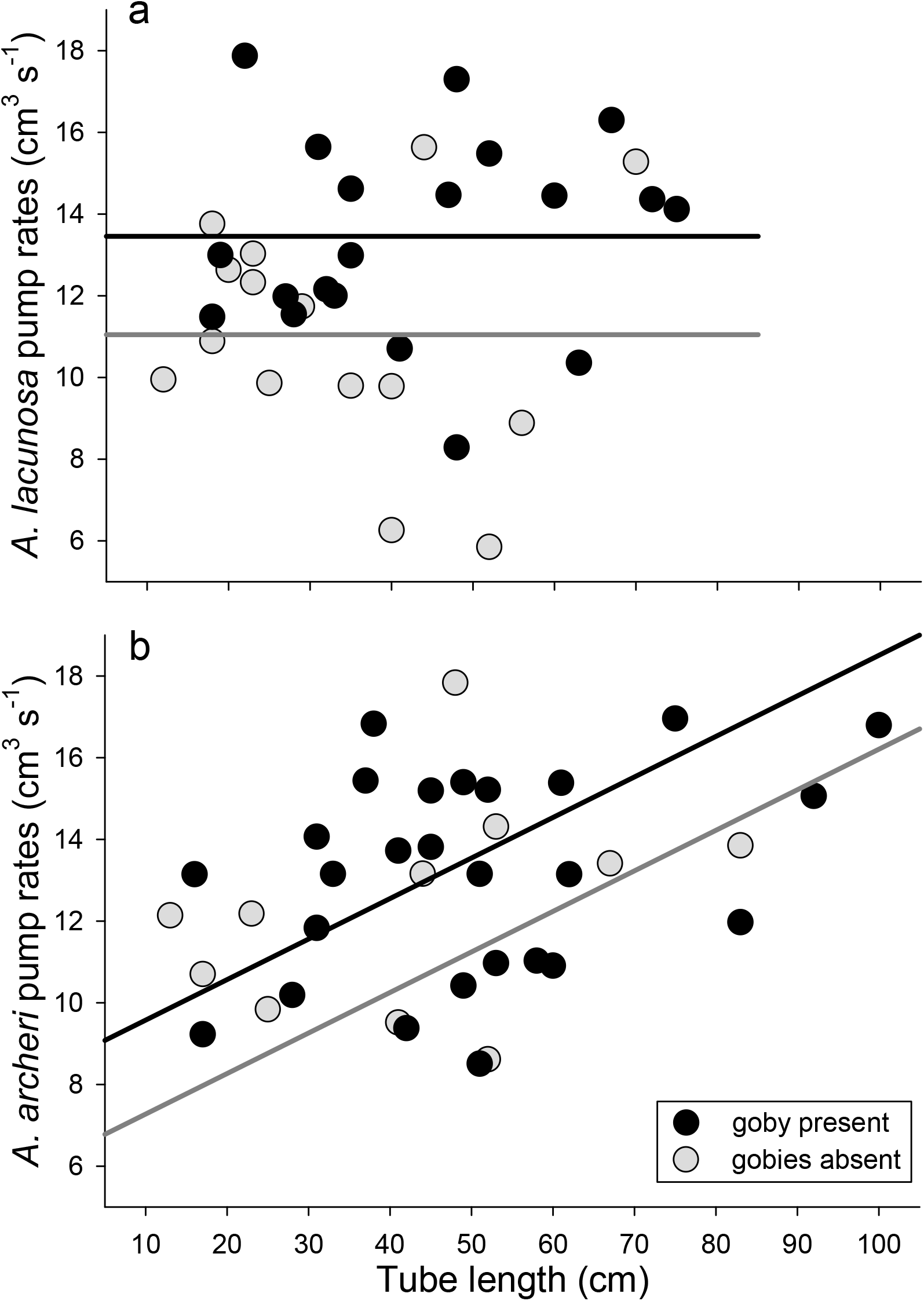
The dependence of average pump rate on tube length and goby presence for a) *Aplysina lacunosa* and b) *Aplysina archeri*. Black symbols and lines indicate a sponge tube with an *Elacatinus horsti*. Gray symbols and lines indicate a sponge tube with no *E. horsti*.

We did not explicitly test parasite reduction as a potential mechanism driving the difference in pump rates; however, other studies have shown that parasitic polychaetes, which live inside sponge tissue and consume choanocytes, have a negative impact on the sponge (Magnino and Gaino 1998). Colin (1975) found that *E. horsti* consume *Haplosyllis* sp., but this conclusion was based on gut content analysis of a small number of fish. In addition, the relationship between *Haplosyllis* sp. and the pump rate of a sponge remains poorly understood (Lattig and Martin 2011). To fully understand the interaction among gobies, sponges, and polychaetes, it is important to examine the relationship between parasite loads and the pump rate of sponges.

The clearing of sediment or algae from sponge tissue has also been suggested as the mechanism of pump rate increases in invertebrate-sponge symbioses because the feeding ability of a sponge decreases when sediment covers their collar cells (Gerrodette and Flechsig 1979; Burns and Bingham 2002). Decreases in pumping rate due to increased sediment has been documented for *A. lacunosa* (Gerrodette and Flechsig 1979). It is possible that *E. horsti* movement and feeding clears sediment from sponge choanocytes in a relationship similar to one seen between brittle-stars and sponges (Hendler 1984). Regardless of the underlying mechanism, we found that goby presence is associated with an improved pumping efficiency which likely increases the feeding ability of the host sponge. These results may apply to other sponge species that host microhabitat dwellers (Tyler and Böhlke 1972; Villamizar and Laughlin 1991; Bell 2008) and is likely to apply to other sponge-dwelling *Elacatinus* species with similar biological and ecological traits.

### Interspecific differences

*Aplysina archeri* tubes were more likely to have an *E. horsti* present than were *A. lacunosa* tubes (z=3.32, p<0.001; Fig. 2). In other southern Caribbean reefs, Villamizar and Laughlin (1991) found a lower frequency of *E. horsti* in *A. lacunosa* tubes compared to *A. archeri* tubes. Differential settlement patterns and persistent recruitment between sponge species and sizes has been recorded for another sponge-dwelling goby species, *Elacatinus lori* (Majoris et al. 2018). The study conducted by Majoris et al. (2018) indicated that habitat preferences in sponge-dwelling gobies are persistent between generations and geographical area, and there is likely a genetic basis for settlement preference. Little is known about the movement of sponge-dwelling gobies post-recruitment (Majoris et al. 2018). Habitat qualities such as food availability and parasite density may influence the settlement of gobies; however, parasite density has not been measured between sponge species or size classes (D’Aloia et al. 2011). Nonetheless, sponge-dwelling cleaning gobies have been shown to preferentially maintain territories on barrel sponges with higher parasite densities (Whiteman and Côté 2004b). The movement of these gobies is critical for understanding the relationships between sponges and gobies.

**Fig. 2.**
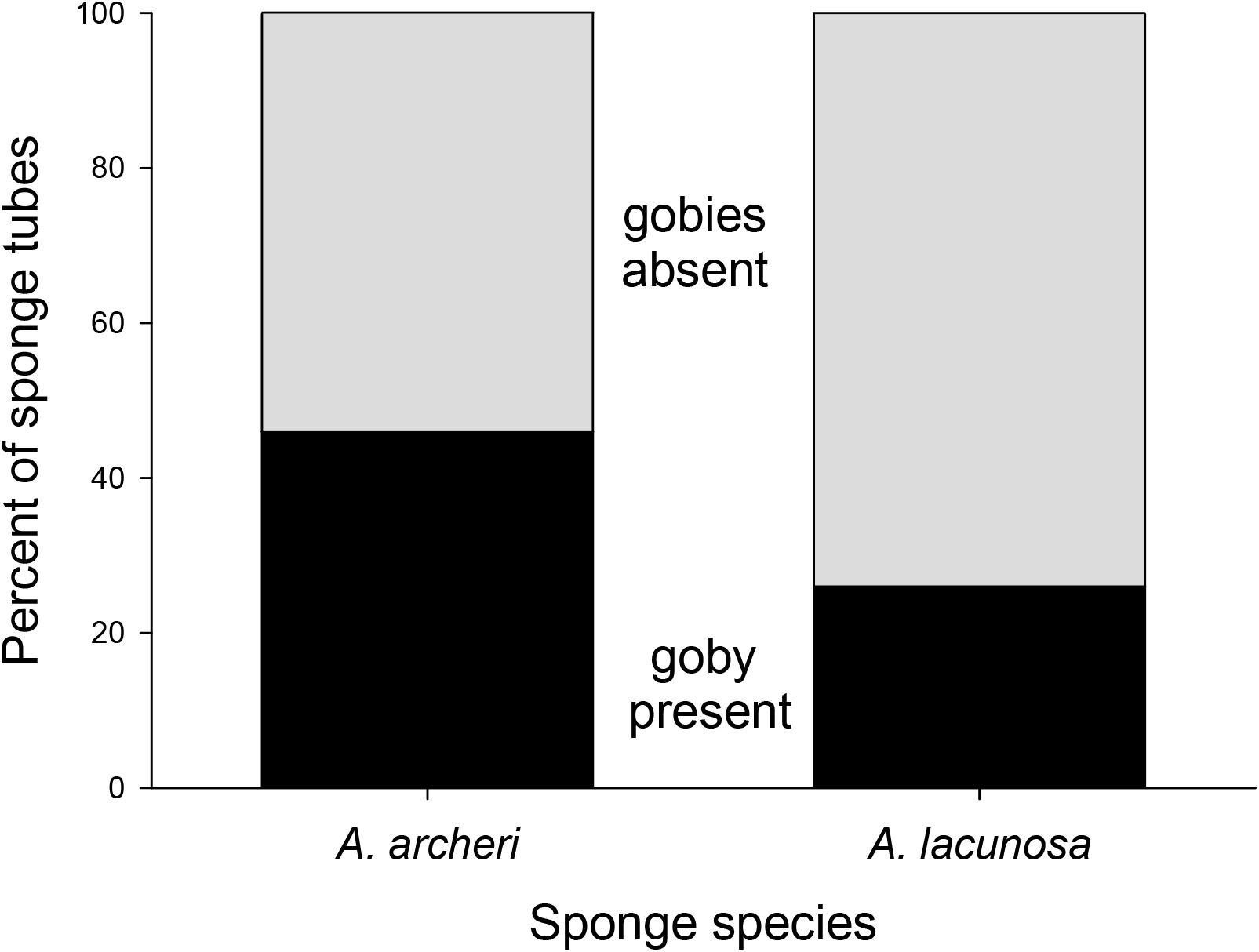
The percent of *Aplysina lacunosa* (n=226) and *Aplysina archeri* (n=85) sponge tubes with an *Elacatinus horsti* (black bar) or with no *E. horsti* (gray bar).

We found that pump rates increased with tube length for *A. archeri* (F_1,34_=24.26, p<0.0001, slope: 0.097 cm^2^ s^-1^) but did not depend on tube length for *A. lacunosa* (F_1,32_=0.09, p=0.7664; Fig 1). Other studies have shown that the pump rates of sponges are higher for longer tubes (Lesser 2006). The pump rates we found here are comparable to those reported previously for sponges of similar size (Gerrodette and Flechsig 1979; Morganti et al. 2019). It is possible that the higher surface area in *A. lacunosa*, due to its corrugation, caused the lack of correlation between the tube length and pump rate we have found for this species (Villamizar and Laughlin 1991). To more clearly understand the effect of tube length on pumping, other factors affecting interspecific differences need to be examined.

### Applications to monitoring reef health

As active suspension-feeders, sponges can impact overall nutrient availability in the water column and, in turn, impact reef health overall (Lesser 2006). As such, sponge health should be considered when evaluating reef health and plans for monitoring reefs. With the relationship between *E. horsti* and *Aplysina* spp. resulting in greater pump rate, goby presence may be an indicator of sponge health. Taking into account these interspecific relationships can help to make reef monitoring more comprehensive (Bell et al. 2017). With current climate trends, accounting for factors that may impact the competitive relationship between corals and sponges may also help scientists to understand and predict future shifts towards or away from sponge-dominated reefs (Bell et al. 2017, 2018).

## Acknowledgements

We would like to thank the Council on International Educational Exchange (CIEE) Research Station Bonaire for support; CIEE station staff and students for dive support and helpful comments; E. Siemann for statistical advice; and S. Brown, J. Majoris, R. Peachey and E. Siemann for comments on this manuscript.

## Data availability

All data are available from the corresponding author upon reasonable request.

